# Impact of Memory T Cells on SARS-COV-2 Vaccine Response in Hematopoietic Stem Cell Transplant

**DOI:** 10.1101/2023.10.26.564259

**Authors:** Jennifer VanOudenhove, Yuxin Liu, Raman Nelakanti, Dongjoo Kim, Emma Busarello, Natalia Tijaro Ovalle, Zhihong Qi, Padmavathi Mamillapalli, Alexa Siddon, Zhiliang Bai, Alfredo Axtmayer, Cheryl Corso, Shalin Kothari, Francine Foss, Iris Isufi, Toma Tebaldi, Lohith Gowda, Rong Fan, Stuart Seropian, Stephanie Halene

## Abstract

During the COVID-19 pandemic, hematopoietic stem cell transplant (HSCT) recipients faced an elevated mortality rate from SARS-CoV-2 infection, ranging between 10-40%. The SARS-CoV-2 mRNA vaccines are important tools in preventing severe disease, yet their efficacy in the post-transplant setting remains unclear, especially in patients subjected to myeloablative chemotherapy and immunosuppression. We evaluated the humoral and adaptive immune responses to the SARS-CoV-2 mRNA vaccination series in 42 HSCT recipients and 5 healthy controls. Peripheral blood mononuclear nuclear cells and serum were prospectively collected before and after each dose of the SARS-CoV-2 vaccine. Post-vaccination responses were assessed by measuring anti-spike IgG and nucleocapsid titers, and antigen specific T cell activity, before and after vaccination. In order to examine mechanisms behind a lack of response, pre-and post-vaccine samples were selected based on humoral and cellular responses for single-cell RNA sequencing with TCR and BCR sequencing. Our observations revealed that while all participants eventually mounted a humoral response, transplant recipients had defects in memory T cell populations that were associated with an absence of T cell response, some of which could be detected pre-vaccination.

## Introduction

Hematopoietic stem cell transplant (HSCT) recipients are at an increased risk for poor outcomes from severe acute respiratory syndrome coronavirus 2 (SARS-CoV-2) infection due to an impaired immune response. A combination of factors contribute to a compromised immune system in HSCT recipients, including history of prior hematologic malignancy, treatment with myeloablative or lymphodepleting chemotherapy, and usage of immunosuppression or further maintenance therapy following transplant (1). Data suggest that all-cause mortality rates from COVID-19 infections in HSCT recipients range between 10-40% (2–5).

Given the increased risk for severe COVID-19 infection and mortality among HSCT patients, proactive infection mitigation is paramount. SARS-CoV-2 mRNA vaccines, including Pfizer’s BNT162b2 and Moderna’s mRNA-1273, were made available under Emergency Use Authorization (EUA) in December 2020 after they showed a 95% and 94.1% efficacy, respectively, in preventing severe symptomatic COVID-19 infection (6, 7). However, these initial trials excluded immunocompromised patients from their enrollment, leading to insufficient data on the efficacy of vaccination in those with immune deficiencies. Despite this uncertainty, vaccination in immunocompromised populations, including HSCT recipients, commenced in 2021. Initial studies were optimistic, showing that antibody response to SARS-CoV-2 vaccination following allogeneic stem cell transplantation ranged from 76-83% (8, 9). In autologous stem cell transplant recipients, Salvini et al. showed that 87% of patients developed a humoral immune response (10). However, in addition to humoral immunity, cellular immune determinants contribute to SARS-CoV-2 infection clearance (11). Given the changes in the cellular immune system following HSCT and exposure to immunosuppressive agents, less is known regarding the cellular immune response following SARS-CoV-2 mRNA vaccination.

Here we report the results of a 22 month long prospective observational study of the immunologic efficacy and comprehensive immune profiling of the SARS-CoV-2 vaccination in 42 hematopoietic stem cell recipients and 5 healthy controls.

## Results

### Patients and Clinical Characteristics

Our study involved 42 transplant patients and 5 healthy controls (HC) as outlined in Table 1. The median age of the HSCT group was 65 years, with ages ranging from 25 to 78 years. Males represented 69.4% (25 of 42) of this group. Out of the 42, 37 were allogeneic transplant recipients (alloSCT), with 81% diagnosed with a myeloid malignancy. The median time from transplant to vaccine (TTV) in alloSCT recipients was 19 months and 12 out of 36 (33.3%) alloSCT patients began their SARS-CoV-2 vaccination less than 12 months after transplant. A significant 56.8% of alloSCT patients had a history of chronic graft versus host disease (cGVHD), and 54.1% were on immunosuppressants (Supplemental Table 1). Among those alloSCT recipients vaccinated more than 12 months after their transplant, 68% (17 of 25) had active cGVHD at the time of their vaccination, compared to 25% of (3/12) alloSCT recipients who started their vaccination series less than 12 months post-transplant. Of the 42 patients, 5 were autologous stem cell transplant recipients (autoSCT). Most of these autoSCT patients had been diagnosed with multiple myeloma with a median TTV of 4 months.

**Table 1.** HSCT Patient Cohort.

|  | **Allogenic (n = 37)** | **Autologous (n = 5)** | **Control (n=5)** |
| --- | --- | --- | --- |
| Age at Vaccine (median, range) | 65 (25-78) | 60 (54-72) | 54 (35-85) |
| Gender (Male) | 25 (69.4%) | 1 (20.0%) | 3 (60%) |
| Hematologic Malignancy |  |  |  |
| AML/MDS/MF | 30 | --- | --- |
| ALL | 4 | --- | --- |
| Lymphoma | 2 | 1 | --- |
| Multiple Myeloma | 1 | 4 | --- |
| Median time to vaccine from transplant (mo) | 19 (7-79) | 4 (4-10) | --- |
| Vaccination ≤12 mo from transplant | 15 (40.5%) | 4 (80%) | --- |
| cGVHD at time of vaccine | 21 (56.8%) | --- | --- |
| On immunosuppression | 20 (54.1%) | --- | --- |
| Lymphopenia (ALC < 1x10<sup>4</sup>/ul) | 9 (24.3%) | 4 (80.0%) | --- |
| Documented COVID prior to vaccine | 3 (8.1%) | 0 | 0 |
| Developed COVID post vaccine | 14 (37.8%) | 1 (20.0%) | --- |

### Serologic Antibody Response

Peripheral blood mononuclear cells (PBMCs) and serum from HSCT patients and healthy controls were prospectively collected before and after each dose of the SARS-CoV-2 vaccine (Figure 1A). Serological testing for SARS-CoV-2 Spike and Nucleocapsid antibodies revealed no significant differences in antibody levels between groups at any vaccination stage (Figure 1B and 1C). However, after the first mRNA vaccine dose, 25% of autoSCT and 13% of alloSCT patients exhibited an effective antibody response (defined as anti-Spike IgG titer > 210 U/mL, based on the value given by the FDA in the EUA for high titer convalescent plasma levels via the same Roche assay), in contrast to 33% of healthy controls. Following a second dose, 75% of autoSCT, and 78.1% of alloSCT patients achieved a robust antibody response. In all cohorts, three doses of the vaccine resulted in a robust response in 92-100% of patients. Comparing alloSCT patients who began vaccination <12 months following transplant to >12 months, there was no significant difference in the rate of achieving a robust antibody response after 2 or 3 doses of the vaccine (Figure 1D and 1E). Among three patients with pre-vaccination samples and previous COVID-19 infections, two tested positive for nucleocapsid antibodies before vaccination; all other patients were negative for Nucleocapsid antibody consistent with low infection rates in patients who followed strict isolation practices. Five of the seven patients who had a post vaccination sample reactive for Nucleocapsid antibodies had reported a COVID-19 infection within the testing window; there were no reported infections within the testing window post initial vaccination that were not caught with nucleocapsid testing.

**Figure 1.**
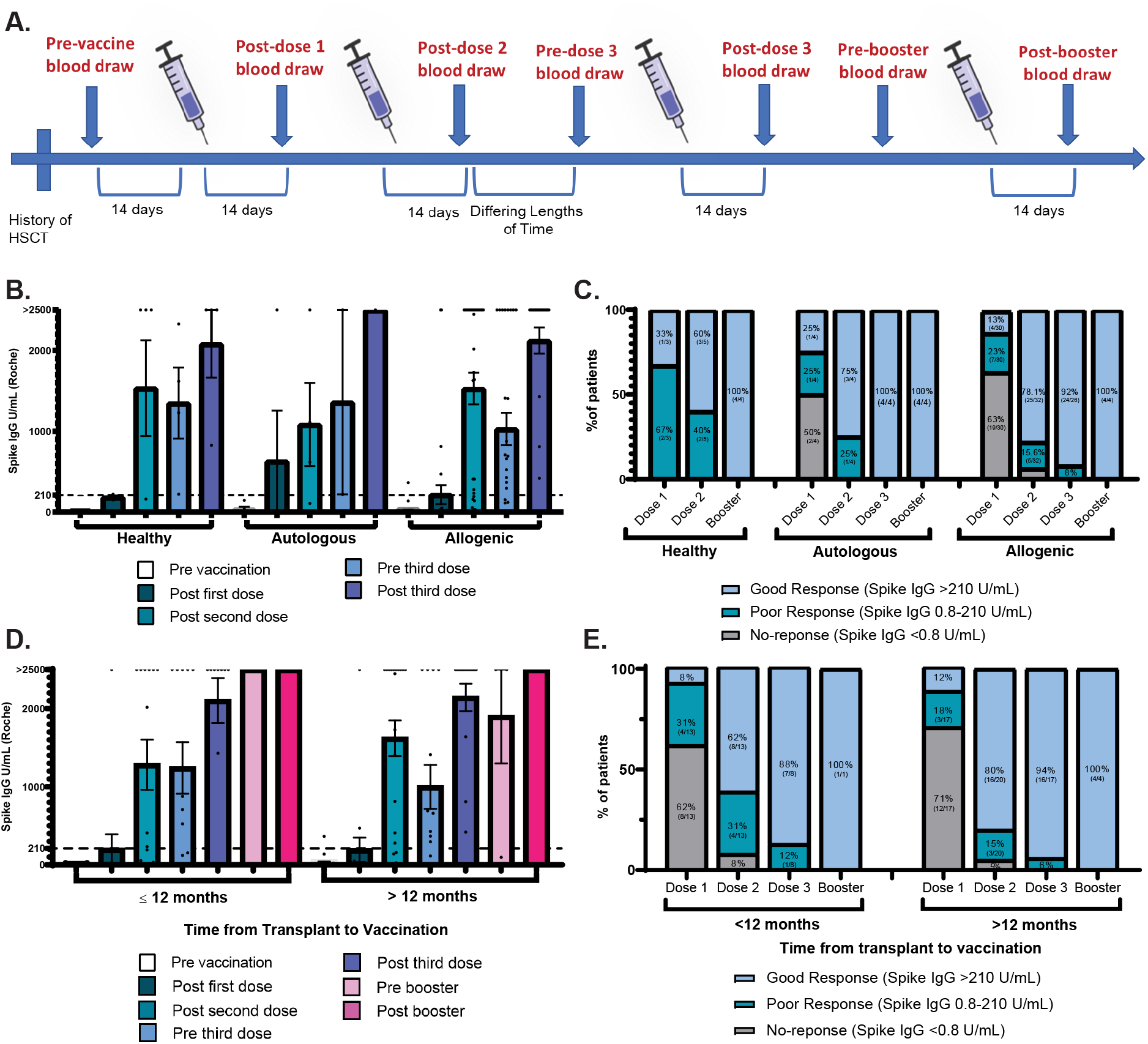
A. Study Design. B. SARS-CoV-2 Spike antibody titers were assessed using the Roche Elecsys anti-SARS-CoV-2 Spike immunoassay. C. Proportion of patients in each cohort who achieved a good, poor, and no antibody response. D. Antibody measurements of allogeneic HSCT recipients who received the SARS-CoV-2 vaccine before and after 12 months following their transplant were plotted by vaccination time points. E. Proportion of allogeneic HSCT recipients (grouped by time of vaccination from transplant) who achieved a good, poor, and no antibody response. Fisher’s exact tests for categorical variables and Mann-Whitney test for continuous variables were performed, with two-sided p-values <0.05 considered statistically significant.

### SARS-CoV-2 Antigen Specific T Cell Activation Assay

Since most patients had an adequate humoral response after the second vaccination dose, we chose to test SARS-CoV-2-specific T cell function via an antigen-specific T cell activation assay on PBMCs of 24 patients (16 alloSCT, 3 autoSCT, 5 control) after the second vaccine dose. The percentage of activated T cells was reported as a percentage of the sample displaying the activation marker above vehicle stimulated cells (Figure 2A and 2B). A positive spike-specific T cell response was defined as having either an increase of 0.1% of double positive TNFα+CD154+ CD4 T cells between pre vaccine and post 2^nd^ dose or an 0.05% increase of double positive IFNγ+TNFα+ CD8 T cells (Figure 2C). In total, 75% of autoSCT and 46.7% of alloSCT patients demonstrated a positive spike-specific T cell response compared to 80% of healthy controls. When comparing the percentage of spike specific CD4 and CD8 T cells before and after 2 doses of vaccination between the 3 cohorts, we observed an increase average percentage of antigen-specific T cells in CD4 T cells in healthy and autoSCT recipients, but not in alloSCT recipients. A modest increase in antigen specific CD8 T cells was seen following 2 doses of vaccine in the healthy and autoSCT cohorts, but not in alloSCT recipients. By univariate analysis of clinical variables and their relationship with a positive T cell response, cGHVD demonstrated an odds ratio of 12.0 that was statistically significant (Figure 3A). A log rank test demonstrated that there was no significant difference in the probability of developing COVID-19 infection among patients who did or did not develop cellular immune response to the vaccine (Figure 3B) and neither log rank test nor uni-or multivariate analysis of clinical variables identified statistically significant relationships with an antibody response (Figure 3C and 3D, Supplemental Table 2).

**Figure 2.**
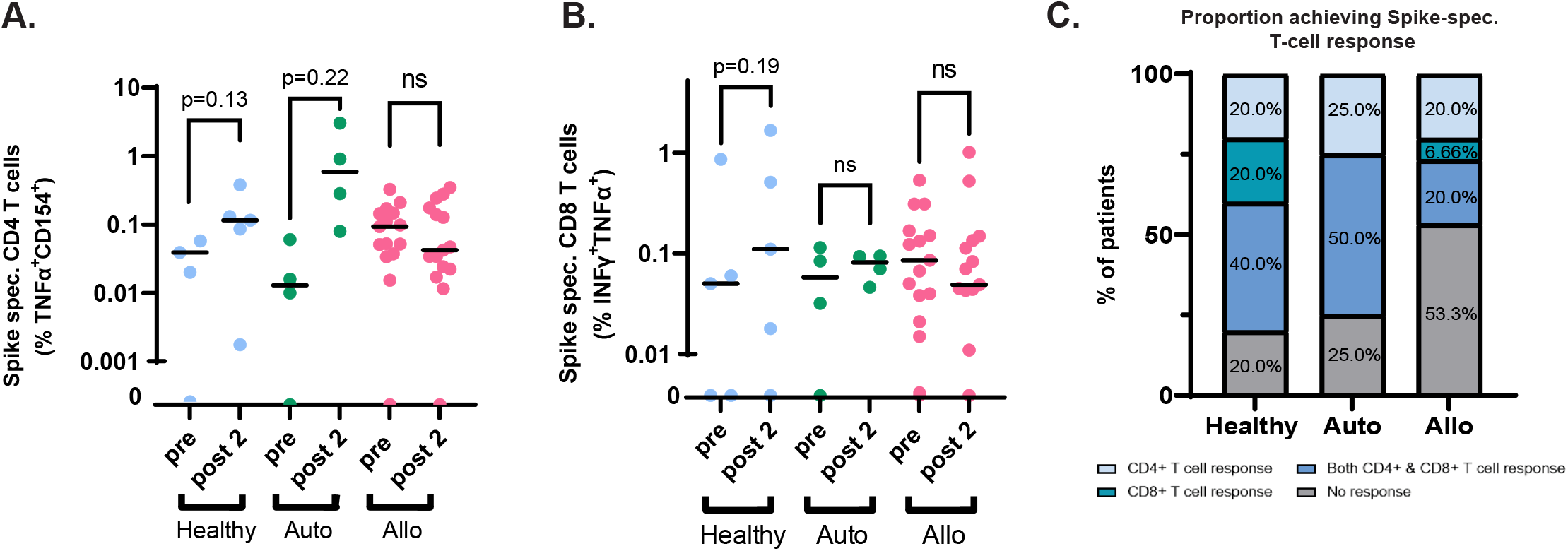
A. Flow Cytometry analysis was performed on PBMCs taken pre-vaccination and following 2nd vaccine dose, cells were stained for intra and extracellular activation markers (CD3, CD4, CD8, CD154, TNF-α, and IFNγ). Percent of spike reactive CD4 cells pre and post dose 2 of the vaccine over vehicle stimulated cells by markers TNFα and CD154. B. Same as A, except for CD8 cells and markers IFNγ and TNF-α. C. Proportion of patients in each cohort who achieved a spike-specific T cell response. Mann-Whitney test for continuous variables were performed, with two-sided p-values <0.05 considered statistically significant (*).

**Figure 3.**
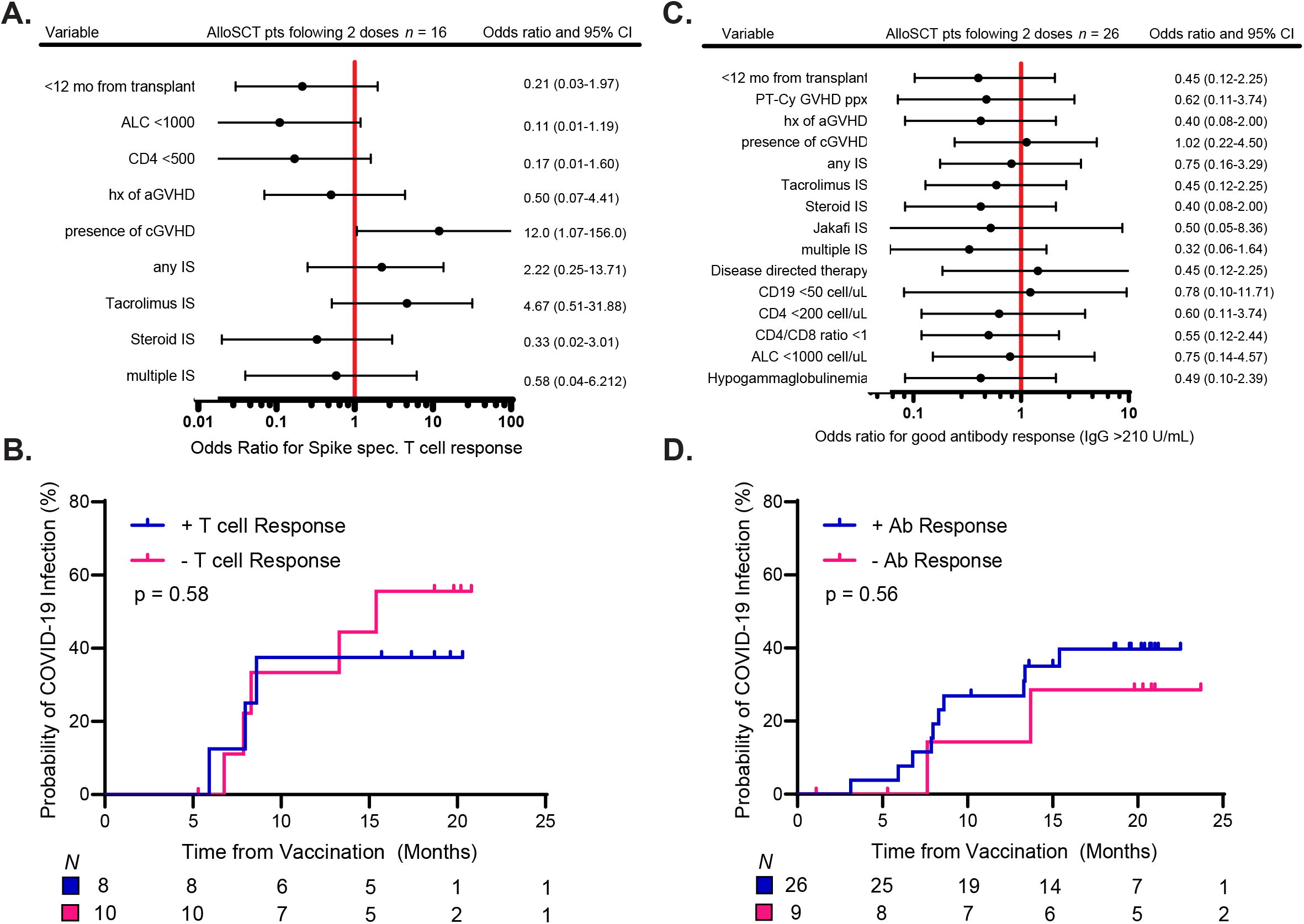
Forest Plots demonstrating clinical variables of interest and their odds ratios of developing a positive T cell response (panel A) or a good (>210 AU/mL) antibody titer (panel C) among allogeneic stem cell recipients. Log Rank Test were performed to test if having a SARS-CoV-2 specific T cell (panel B) or a significant antibody (panel D) response resulted in a difference in probability of COVID infection.

### Immune profiling

#### Clinical Flow Cytometry for Immunodeficiency

A thorough evaluation of the cellular immune repertoire was conducted by flow cytometry for 39 out of 42 HSCT recipients prior to vaccination. Immune cell subset analysis revealed that compared to autoSCT recipients, alloSCT recipients had a significantly higher absolute CD19 B cell count (allo: 139.7 cell/µL, auto: 22.5 cells/µL; p <0.01) and increased relative frequencies of memory B-cell (allo: 0.55%, auto: 0.2%; p=0.02), class switched B-cell (allo: 0.73%, auto 0.2%; p <0.01) and gamma-delta T cells (allo: 2.3%, auto 0.8%; p <0.01) (Figure 4A). Furthermore, alloSCT recipients demonstrated lower levels of activated T cells, as evidenced by the ratio of HLA-DR+ to HLA-DR-T cells (allo: 0.38 versus auto: 0.77, p = 0.06). We then examined whether any immune cell subset present prior to vaccination could predict antibody responses after two doses of the SARS-CoV-2 mRNA vaccine (Figure 4B). Transplant recipients with poor antibody response, identified by a spike IgG titer of less than 210 U/mL, had a significantly higher percentage of CD45RA naïve T cells at baseline compared to those with a good antibody response (poor responders: 30%, good responders: 19.93%; p = 0.02). Additionally, an evaluation of the relationship between baseline immune subsets and antigen specific T cell response was conducted (Figure 4C). Patients who did not develop a SARS-CoV-2 spike-antigen specific T cell response had a lower absolute lymphocyte count (ALC) (responders: 1.7*10 ∧ 4 cells/µL, non-responders: 0.8*10∧4 cells/µL; p < 0.01) with decreased counts of CD4 T cells (responders: 442 cells/µL, non-responders: 335 cells/µL; p = 0.09) and CD8 T cells (responders: 568 cells/µL, non-responders: 159 cells/µL; p < 0.01). The frequency of CD45RO+ memory T cells trended lower but without statistical significance (responders: 30.6%, non-responders 19.62%; p = 0.14).

**Figure 4.**
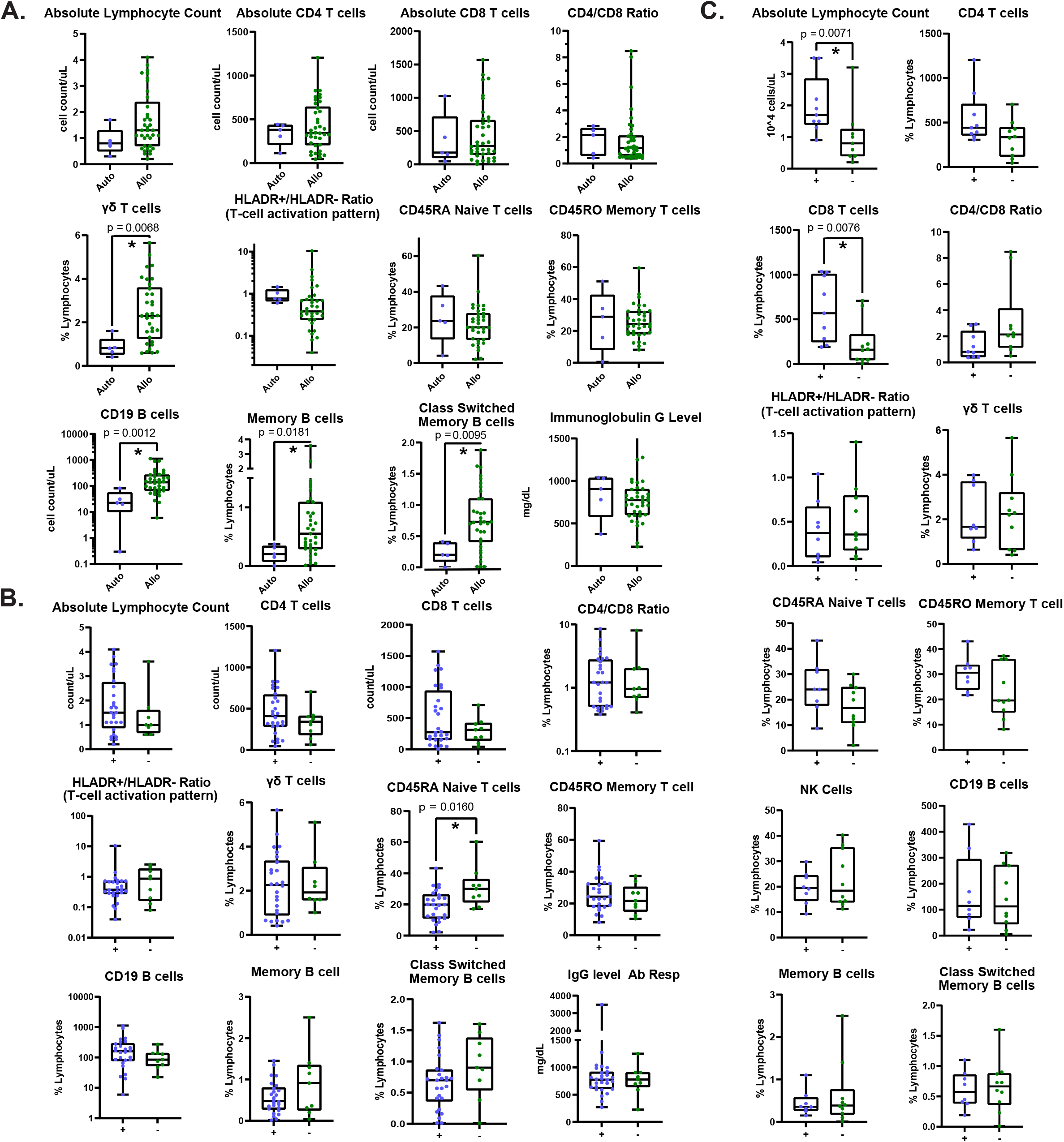
Clinical Pre-vaccination Immunodeficiency Flow Panel. A. Results presented by transplant type. B. Results presented by positive or negative significant antibody response. C Results presented by positive or negative SARS-CoV-2 specific T cell stimulation assay response. Mann-Whitney test for continuous variables were performed, with two-sided p-values <0.05 considered statistically significant (*).

#### CyTOF

Mass Cytometry was carried out on PBMCs prior to vaccination and after the first and second doses in 8 alloSCT patients, 2 autoSCT patients, and 2 healthy individuals. Among this group, 58.8% of patients developed a good antibody response and 72.3% developed a positive T cell response after two vaccine doses. No significant variations in immune cell subsets were observed between the times of vaccination. Immune cell subsets at baseline were examined among the three different groups before vaccination (Figure 5A). There were fewer central memory CD8 T cells in alloSCT patients compared to healthy controls (healthy: 3.01%, alloSCT, 0.87%; p = 0.04) and autoSCT (autoSCT: 2.7%, p = 0.11). In contrast, plasmablasts were more numerous in alloSCT cohort compared to healthy controls (healthy: 0.07%, alloSCT: 0.26%; p =0.09). Analyzing the groups based on their ability to produce antibodies revealed that those with poor antibody response had fewer plasmablasts (responders: 0.12%, non-responders: 0.31%; p = 0.19), and higher percentage of terminal effector CD4 T cells (responders: 47.3%, non-responders: 53.0%; p = 0.09) (Figure 5B). Lastly, patients who did not display an antigen-specific T cell reaction had reduced percentages of CD3 T cells (responder: 6.3%, non-responder: 63.9%; p = 0.07), central memory CD4 T cells (responder: 2.1%, non-responder 0%, p = 0.26), and central memory CD8 T cells (responder 1.8%, non-responder 1.3% p = 0.6) (Figure 5C).

**Figure 5.**
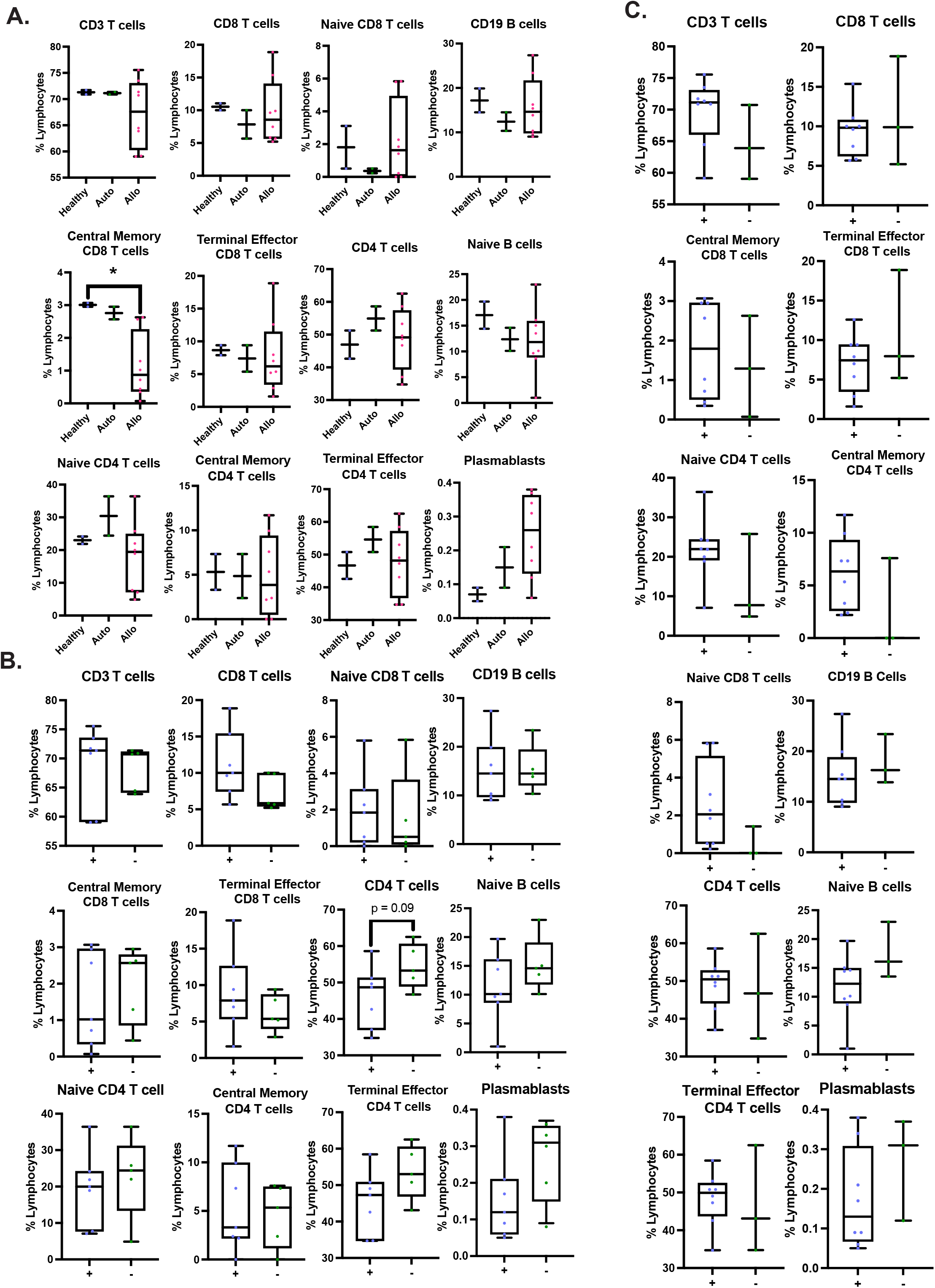
CYTOF. A. Cell types presented as percentage of lymphocytes separated by transplant type. B. Cell types presented as percentage of lymphocytes separated by positive or negative significant antibody response. C. Cell types presented as percentage of lymphocytes separated presented by positive or negative SARS-CoV-2 specific T cell stimulation assay response. Mann-Whitney test for continuous variables were performed, with two-sided p-values <0.05 considered statistically significant (*).

#### Single cell RNA-sequencing

To better understand predictors of cellular responses to SARS-CoV-2 mRNA vaccines in SCT recipients, we performed 10X single cell RNA-seq (scRNA-seq) combined with T Cell Receptor (TCR) and B Cell Receptor (BCR) sequencing on pre-and post-vaccination (2^nd^ dose) samples, as well as post-infection samples when available, from 4 alloSCT recipients and 3 healthy controls. These subjects were chosen to include both good and poor humoral and T cell responders to vaccination. The clinical characteristics of the patients selected for single cell RNA seq are provided in Supplement Figure 1A. From the seven individuals sequenced, 117,492 cells passed quality control to exclude doublets, cells with high mitochondrial reads, and feature counts below or above what would be expected for a single cell (Supplemental Figure 1B-1C). High quality cells were clustered based on their gene expression, and each cluster was assigned a cell type identity based on reference annotation dataset (“MonacoImmuneData”) (Supplemental Figure 1D)(12). The presence or absence of a T or B clonotype in the TCR and BCR cell sequencing was used to further support the cluster identification. Next, T and B cells were selected and re-clustered for subset analyses (Figure 6A and 6B). The expression of maker genes *CD3E*, *CD4*, *CD79A*, and *CD8A*, as well as the top marker genes in heatmap form, are shown to support the cluster identities (Figure 6A-C).

**Figure 6.**
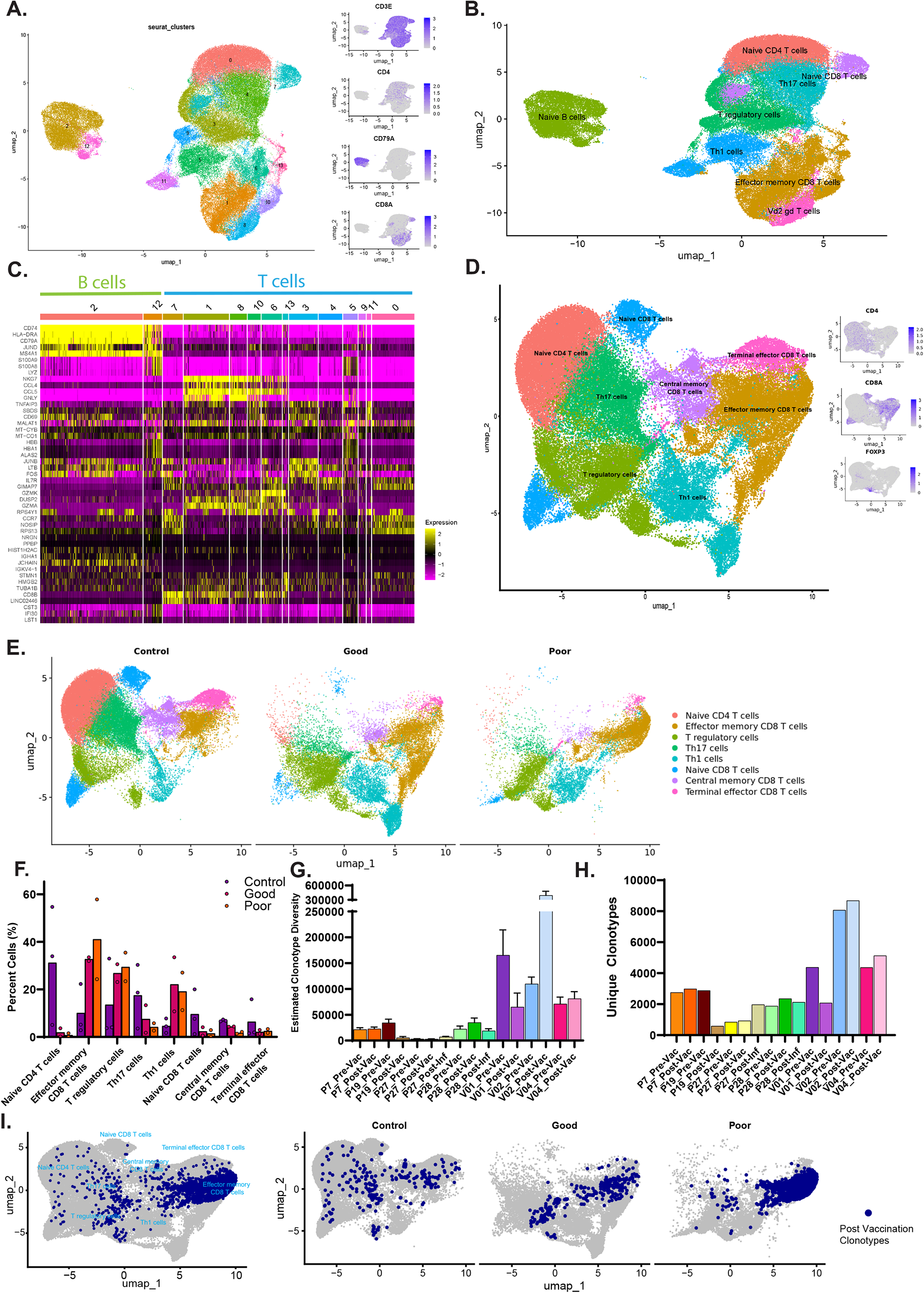
Single cell RNA sequencing of patient samples and TCR clonotype analysis. A. UMAP projection of the 105,800 lymphocytes with numbered Seurat clusters labeled. Representative genes identifying cell types within the UMAP (right) including *CD3E*, *CD4*, *CD79A*, and *CD8A*. B. Seurat clusters with identities annotated using singleR and the Monaco dataset. C. Top gene markers identified for each cluster plotted in a heatmap. D. Data was filtered to include only T cell clusters and excluded two post-COVID-19 infection samples. UMAP projection of the T cells re-clustered and re-annotated as before with markers *CD4*, *CD8*, and *FOXP3*. E. Cells from the UMAP projection in D were split by the individual’s T cell response and colored by cluster annotation. F. Quantitation of E, where the percentage of each individual’s cells in each cluster was calculated (Healthy n=3, Good n=2, Poor n=2) and plotted on a bar graph. G. TCR repertoire diversity using the Chao1 estimator, the calculated value is plotted as a bar graph displaying the calculated 95% confidence interval. H. A bar graph displaying the number of unique clonotypes in each sample sequenced. I. Top clonotypes found to be unique to post vaccination when compared to pre vaccination for each individual were mapped back to the cells in the T cell UMAP both in aggregate (left) and the UMAP with cells divided by response to antigen specific T cell assay cohort (right).

The T cells were then analyzed separately, re-clustered, and re-annotated using fine labeling from the same reference annotation (Figure 6D). T cell markers *CD4* and *CD8*, as well as regulatory T cell marker *FOXP3*, expression within the cells was again shown to support cluster annotation, along with a heat map of the top marker genes (Figure 6D and Supplemental Figure 1E). To determine whether changes in T cell frequencies were evident in good versus poor responders compared to healthy controls (Figure 6E), we calculated the percentage of cells in each cluster for each individual sequenced (Figure 6F). All alloSCT recipients displayed decreased naïve CD4+ T cells, naïve CD8+ T cells, Th17 cells, and terminal effector CD8 T cells compared to healthy controls. AlloSCT recipients with a good T cell response, like healthy controls, had more memory CD8+ T cells compared to alloSCT recipients with a poor T cell response. Intriguingly, this scRNA-seq data are supported by the CYTOF analysis performed in a larger number of our patients. AlloSCT recipients who demonstrated either a good or poor T cell response had more Th1 and T regulatory cells than healthy individuals. Poor responders had the highest level of effector memory CD8+ T cells when compared to good responders and healthy controls. Gene set enrichment analysis comparing T cell gene expression post vaccination in good versus poor T cell responders revealed enrichment for a T memory cell signature in good responders (Supplemental Figure 1F, and Supplemental Table 3) emphasizing the differentially expressed T memory marker genes *CCR7*, *LEF1*, *IL7R*, and *TCF7*.

Each sample’s T cell repertoire and diversity were further examined by using the corresponding data from the TCR sequencing. The Chao1 estimator function, a nonparametric asymptotic estimator of species richness, was utilized to calculate the clonal diversity on the amino acid level for each individual at each vaccination timepoint (Figure 6G). Then we determined the total number of unique clonotypes for each individual at each time point (Figure 6H). First, as expected, healthy donors had more clonal diversity in comparison to HSCT patients. Specifically, individual P27, followed by P19, had the lowest post vaccination clonotype diversity and unique clonotypes. Interestingly, individual P27 had both poor antibody response and poor antigen specific T cell response, while P19 was able to mount an antibody response but not an antigen specific T cell response. Individuals P7 and P28, who had both good antibody and T cell responses, had higher clonotype diversity and number of unique clonotypes than the poorer responders, but not as high as the control healthy individuals.

Finally, the top clonotypes for each individual were queried for uniqueness between the pre and post vaccination time points, as clonotypes that appear post vaccination could be related to the vaccination event, and 256 unique clonotypes were identified (Supplemental Table 4). This list was then annotated using a database, TCRMatch, to determine the clonotypes’ potential predicted targets (13). Of the 109 out of 256 clonotypes that matched with an entry in the annotation database, 67 were predicted to target a portion of SARS. The unique post vaccination clonotypes were then mapped back to the cells in the T cell UMAP (Figure 6I). When overlayed onto the T cell UMAP there was a noticeable concentration of cells in the effector memory CD8+ T cells cluster. To determine whether there was any bias in which patients these cells were from, we split the overlay according to T cell response. For the control individuals the post vaccination clonotypes were evenly distributed across the different cell types. However, with the HSCT patients, the distribution of post vaccination clonotypes was much more restricted. Between the good and poor T cell responders, there was an even more predominate concentration of post vaccination clonotypes in the effector memory CD8+ T cell cluster of the poor responders. This suggests a defects in the T cells functional differentiation dynamics, potentially preventing cells from reaching the CD8+ central memory or terminal effector state (Supplemental Figure 1G).

Parallel investigations executed on the B cells mirrored the T cell analyses. However, there were no identified trends in B cell populations or antibody response. This was expected given the universal humoral response in healthy controls and our alloSCT recipients (Supplemental Figures 1H-1J).

### Clinical Outcomes

As of the last data update on November 26, 2022, 14 out of 37 alloSCT recipients (37.8%) and 1 out of 5 autoSCT patients (20%) had developed a COVID-19 infection. Among the 15 affected patients, two were admitted to the general medicine floor in the hospital, with no instances of ICU admissions or COVID-19 related fatalities. Three individuals passed away post-vaccination, but their deaths were not related to COVID-19 infection or vaccination complications. A Kaplan Meier analysis revealed no significant disparity in COVID-19 infection rates between patients with and without a strong antibody response, as visualized in Figure 3C. Importantly, no adverse reactions to the vaccine were reported.

## Discussion

Our findings provide a detailed analysis of the humoral and cellular immune response in allogeneic stem cell transplant recipients to the initial SARS-CoV-2 mRNA vaccines and offer insights that may influence clinical decision making for vaccination in this population. Our studies highlight a diminished antibody response in transplant recipients relative to healthy controls during the initial vaccine doses, most markedly after the first dose. Nevertheless, the introduction of a third standard dose, accompanied by subsequent boosters, markedly uplifts the proportion of good responders, aligning it with the response seen in healthy individuals.

Vaccination efficacy is often compromised in allogeneic recipients early after transplantation and routine revaccination is typically postponed at least 10-12 months post transplantation. We have shown that selected patients can respond at an early timepoint post-transplant to SARS-CoV-2 vaccination. This contrasts with other studies that have demonstrated early vaccination from time of transplant (<12 months) and immunosuppression have been associated with poor serologic responses to vaccination (14, 15). Instead, in our cohort patients vaccinated on immunosuppression or that were <12 months following transplant achieved similar antibody responses to their counterparts. However, it should be noted that in our cohort, patients selected for vaccination less than 12 months from transplant were purposefully those patients off or almost off immunosuppression and without GVHD, which could be contributing to the lack of association between time from transplant and antibody response.

Beyond the anti-spike antibody response, the SARS-CoV-2 mRNA vaccine has also proven effective in eliciting potent T cell reactions in healthy subjects. Vaccine-induced CD8+ T effector cells may be detected as early as 10-days after the primary dose (16). Our immune profiling indicates that HSCT patients who did not exhibit a spike-specific T cell response had reduced lymphocyte counts, including both CD4 and CD8 T cells, and fewer CD45RO+ memory T cells compared to those who did exhibit a spike-specific T cell response. This is further corroborated by our scRNA-seq data that demonstrates a larger CD8 central memory T cell compartment and higher memory T cell gene expression in the healthy control and good T cell responding alloSCT patients compared to the non-responders. The central memory T cell clusters in the post-vaccine samples appear stable from their pre-vaccine timepoint, which is consistent with prior data showing that a booster dose conserves the memory T cell pool (17). Additionally, we found an accumulation of post vaccination specific clonotypes in the effector memory CD8+ T cell cluster is most pronounced in the poor responding cohort, suggesting an additional defect in the functional differentiation of the T cell memory compartment.

Our study offers a comprehensive evaluation of the immune status of transplant recipients before and after vaccination forging a path for subsequent scrutiny into immune reconstitution and vaccine responses in this demographic. Although specialized analytical techniques, such as mass cytometry and single-cell sequencing, are not available in routine clinical care, we have demonstrated that there is concordance between clinical flow cytometry, mass cytometry, and single-cell sequencing. Therefore, more accessible clinical evaluations, such as absolute lymphocyte count and routine lymphocyte subset flow cytometry delineating CD4+ and CD8+ T cells, could serve as proxy indicators for a stem cell transplant recipient’s likelihood to develop a humoral and cellular immune response to the SARS-CoV-2 vaccine.

We recognize the limitations inherent in our study, particularly its modest sample size and diverse patient group. It’s crucial to understand that the majority of COVID-19 cases in our cohort arose during the Omicron variant surge, where even vaccinated healthy individuals faced high breakthrough infection rates due to the original vaccination series targeting the Alpha variant.

## Methods

### Sample Collection and Patient Population

Peripheral blood was collected in red top tubes containing no anti-coagulant, allowed to clot, then centrifuged at 1500g for 10 minutes at 4°C, the supernatant was then collected as serum. Peripheral blood samples collected in lavender EDTA coated tubes were layered over Ficoll and separated by density gradient centrifugation at 1800RPM for 20 minutes with the brake off. Peripheral blood mononuclear cells (PBMNCs) were collected from the buffy coat layer, washed with PBS, counted, and then cryopreserved in FBS with 10% DMSO. Patient clinical data including diagnosis, type and timing of transplant, immunosuppression use, history of chronic graft versus host disease (cGVHD), and COVID-19 infection before or after vaccination were collected from the electronic medical record. The samples and majority of data used in this paper were collected between January 2021 and November 2022.

### Serologic Antibody Measurement

Spike and Nucleocapsid antibody titers were assessed using the Roche Elecsys anti-SARS-CoV-2 Spike immunoassay (Spike), Roche Elecsys® SARS-CoV-2 Antigen assay (Nucleocapsid) and a novel microfluidic serologic device (18). A robust spike IgG antibody response was defined as >210 AU/mL for the Roche assay and >38 BAU/mL in our novel device. The Roche Nucleocapsid assay is read out with an Index value that is then categorized as Reactive for an Index value >1.

### T cell Activation Assay

Antigen-specific T cell stimulation was performed on PBMCs taken pre-vaccination and following second vaccine dose with the Miltenyi Biotec SARs-CoV-2 Prot_S T Cell Analysis Kit and Peptivator® Prot-S Peptide Pool. The peptide pool contains the immunodominant sequences to the Wuhan SARS-CoV-2 spike protein (aa 304-338, 421-475, 492-519, 683-707, 741-770, 785-802, and 885 – 1273). Cells were thawed from liquid nitrogen storage and rested overnight at 37°C. PBMCs were plated with 1*10∧6 cells per well and then stimulated according to the manufacturers protocol for 2 hours before addition of Brefeldin A to inhibit exocytosis of activation signaling proteins, and then stimulation continued for an additional 4 hours. Cells were stained for intra and extracellular activation markers (TNFα, IFNγ, CD154/CD40L), as well as T cell markers (CD3, CD4, CD8). Flow cytometry analysis was performed using a BD Symphony Cell Analyzer and analyzed using Flow Jo (v10).

### Immune Profiling

A clinical flow multiparametric flow cytometry designed to evaluate immunodeficiency syndromes was performed by YNHH Department of Laboratory Medicine on whole blood samples of our transplant cohort prior to first vaccine dose. Biobanked PBMCs before and after first and second of the SARS-CoV-2 vaccine were stained with the Fluidigm Maxpar Direct Immune Profiling assay (containing 30 markers that allows discrimination of 37 cell types) and mass cytometry was performed with data acquisition on the CyTOF Helios (Fluidigm/Standard BioTools). Population calling was performed using the Maxpar Pathsetter program (Fluidigm/Standard BioTools).

### Single-cell RNA-seq

Thawed PBMCs from pre-vaccination, following 2^nd^ vaccine dose, and following COVID-19 infection underwent CD19 and CD3 enrichment utilizing MicroBeads and LS Columns on the QuadroMACS™ Separator (Miltenyi Biotec). Single-cell suspensions of ∼1,200 cells/µl in PBS were prepared for scRNA-seq using 10X Chromium Next GEM Single Cell 5’ Kit v2 also using the Chromium Single Cell Human TCR and BCR Amplification Kits (10x Genomics, Pleasanton, CA). Libraries were constructed per the manufacturers protocol. Libraries were sequenced on an Illumina NovaSeq 6000 with 150bp paired end reads to ∼240 million reads for the gene expression libraries and 80 million reads each for the TCR and BCR libraries.

### Single-cell RNA-seq data processing

The demultiplexed fastq files were run through the Cell Ranger multi pipeline (10X Genomics) to generate the gene expression count matrix, T cell V(D)J and B-cell V(D)J analysis files. The data files from each sample were then combined and normalized by running through the Cell Ranger aggr pipeline. These aggregated data matrices were then used for downstream analyses.

Gene expression data analysis was performed using R (v4.3) and the Seurat package (v5) following a standard workflow (19). In brief, the aggregated filtered feature matrix with 118,950 cells was loaded into a Seurat data object. Poor quality cells with greater than 20% mitochondrial gene expression were filtered out, as well as suspected doublets and empty reactions using nFeature < 200 or nFeature >= 4000. Using the V(D)J metadata, cells with both T-and B-cell clonotypes were considered doublets and removed (966 cells).

After filtering, 117,492 cells remained for downstream analysis. The data were normalized and scaled using SCTransform using default parameters with an additional parameter to regress out percent mitochondrial gene expression. After running PCA with default conditions for dimensionality reduction, cells were visualized using UMAP and clusters were identified using the default FindClusters method and a resolution of 0.4. Cluster identities were predicted using SingleR based on a reference immune cell data set “MonacoImmuneData” (12, 20). Gene markers for each cluster further confirmed the cluster identities. Clusters containing monocytes, dendritic cells, and other non-lymphocytes were removed, after which 105,800 lymphocytes remained.

To avoid the influence of high-expressing B-and T cell receptor clonotypes in visualization and clustering, these genes were removed from the lymphocyte dataset count matrix using the patterns “TR[ABDG][VJC]” or “IG[HJKL]” (21). The data were renormalized, visualized with UMAP, and clustered as above. Cluster identities were again predicted using SingleR, as above. Then, the data object was subset into separate T cell and B cell analyses.

The two samples taken after COVID positivity were excluded for both T and B cell analyses and only pre-and post-vaccination analyses were conducted. The T cell dataset had 37,988 and 37,469 pre-and post-vaccination cells respectively. The B cell dataset had 13,049 cells total.

Cluster identities were predicted using SingleR, as above. For differential gene expression between T cell responders (Good versus Poor), the object was first subset into separate analyses for pre-and post-vaccination. Gene set enrichment was performed using Enrichr (22–25). TCR and BCR clonotypes were added as metadata to the Seurat objects above using the filtered contig annotations from the VDJ sequencing. Only the primary clonotype was kept per barcode. The immunarch library (v1.0) was used to generate all clonotype summary data as well as identify top clones per patient sample (26). To identify unique post vaccination clonotypes the top clonotypes were compared for each patient pre and post vaccination. This list was then annotated using the TCRMatch tool to identify epitopes targeted by the TCR clonotypes (13). Cells mapped to this list of clonotypes were then selected and highlighted in the T cell UMAP divided by T cell response. Monocle3 was used to perform trajectory analysis on the T cell UMAP (27–29).

### Study approval

The study was conducted according to the Declaration of Helsinki and was approved by the Institutional Review Board at Yale University (HIC#1401013259).

### Statistical Analysis

Descriptive statistics were performed. Fisher’s exact tests for categorical variables and Mann-Whitney test for continuous variables were performed, with two-sided p-values <0.05 considered statistically significant. Kaplan Meier Survival analysis and log rank tests were conducted in GraphPad Prism v9.2.0

### Data availability

Values for all data points found in graphs can be found in the Supporting Data Values file, and all the raw and processed single cell RNA sequencing TCR and BCR sequencing data in this study can be accessed in the NCBI Gene Expression Omnibus (GEO) database under the accession number GSEXXXXX.

## Author Contributions

J.V and Y. Liu, under advisement from S.H, R.F. and S.S, designed and executed the research experiments, acquired and analyzed data, and co-wrote the manuscript. The order of co–first authors JV and YL was determined by management of the project. A.A, C.C, S.K, L.G, and S.S assisted with sample acquisition. P.M helped with sample processing. D.K and Z.Q helped with data acquisition. R.N, E.B, T.T, Z.B, and A.S contributed to data analysis. N.T.O helped with manuscript preparation. The manuscript was edited and reviewed by all authors.

## Acknowledgments

We would like to acknowledge the assistance of the entire hematopoietic transplant team at YNHH, as without them this work would not be possible. Additionally, we would like to thank Rocco Carbone of the Rapid Case Ascertainment (RCA) Shared Resource of the Yale Cancer Center (YCC) for his assistance with monitoring patient blood collections. We thank Yale Flow Cytometry for their assistance with use of the BD Symphony. The Core is supported in part by an NCI Cancer Center Support Grant # NIH P30 CA016359. The BD Symphony was funded by shared instrument grant # NIH S10 OD026996. We would like to thank Guilin Wang and the YCGA sequencing core, as research reported in this publication used their services which are supported by the National Institute of General Medical Sciences of the National Institutes of Health under Award Number 1S10OD030363-01A1. This work was also supported by an NCI grant awarded to S.H and R.F (U01 CA260507) and funding from t*he Frederick A. DeLuca Foundation*.

