## Supplemental Table 1 for "Impact of Memory T Cells on SARS-COV-2 Vaccine Response in Hematopoietic Stem Cell Transplant"

**Supplemental Table 1. Immunosuppression of Allogenic Patients**

| Characteristic | Overall (n=37) | |
|---|---|---|
| **Immunosuppression (among allogenic)** | | |
| Any Immunosuppression | 20 | (54.1) |
| Steroids | 12 | (23.4) |
| Tacrolimus | 16 | (43.2) |
| JAK2 inhibitor | 3 | (8.1) |
| Photopheresis | 3 | (8.1) |
| Multiple immunosuppressants | 9 | (24.3) |
| **Therapy (within 3 mo of vaccination)** | | |
| Total | 11 | (25.6) |
| Revlimid | 3 | (7.0) |
| Daratumumab | 1 | (2.3) |
| Carfilzomib | 1 | (2.3) |
| Gilteritinib | 3 | (7.0) |
| Venetoclax | 2 | (4.7) |
| Azacitidine | 1 | (2.3) |
| Ibrutinib | 1 | (2.3) |
| Receiving IVIG | 4 | (9.3) |
