## Supplemental Table 2 for "Impact of Memory T Cells on SARS-COV-2 Vaccine Response in Hematopoietic Stem Cell Transplant"

**Supplemental Table 2. Binary Logistic Regression analysis of variables influencing achievement of good antibody response (Spike IgG > 210 U/mL).**

| | **p-value** | **Odds Ratio (95% Confidence Interval)** |
|---|---|---|
| Age | 0.467 | 0.950 (0.826-1.091) |
| Post-Transplant Cyclophosphamide | 0.428 | 5.703 (0.077, 422.631) |
| Time to Vaccination* <12 mo | 0.848 | 0.720 (0.025, 20.628) |
| History of acute GVHD | 0.337 | 0.244 (0.014, 4.357) |
| cGVHD at time of vaccine | 0.779 | 2.086 (0.031, 59.897) |
| Immunosuppression at time of vaccine | 0.900 | 1.363 (0.011, 175.402) |
| ALC < 1000 | 0.648 | 0.383 (0.006, 23.473) |
| CD4 T cell count < 200 | 0.950 | 1.123 (0.031, 41.141) |
| CD19 B cell count < 50 | 0.275 | 1.011 (0.991, 1.032) |
| CD45RA T cells | 0.171 | 0.916 (0.808, 1.039) |
| Hypogammaglobulinemia | 0.872 | 1.366 (0.031, 59.897) |

*Time to vaccination: time from transplant to first vaccine. ALC: absolute lymphocyte count; CD4
