## Supplemental Figure 1 for "Impact of Memory T Cells on SARS-COV-2 Vaccine Response in Hematopoietic Stem Cell Transplant"

**A.**

| ID | Gender | Age | Hematologic Malignancy History | TTV (mo) | Allogenic Transplant Type | Post Transplant Cytoxan | aGVHD history | cGVHD Present | Immuno-suppression | GVHD flare after vaccine | GVHD Treatment | Abs CD4+ (count/ul) | Abs CD8+ (count/ul) | Abs CD19+ (count/ul) | Absolute Lymphocyte count (x1000/uL) | Antibody Response/ T Cell Response | Vaccination Brand |
|---|---|---|---|---|---|---|---|---|---|---|---|---|---|---|---|---|---|
| P7 | F | 61 | Therapy related AML | 12 | MRD | No | No | Yes | NA | Yes | Steroid, Tacrolimus | 590.4 | 207 | 428.4 | 1.5 | Good/Good | Pfizer |
| P19 | F | 72 | Secondary AML | 7 | Haplo | Yes | No | No | NA | No | NA | 499.8 | 222.6 | 278.6 | 1.4 | Good/Poor | Pfizer |
| P27 | F | 77 | MDS | 7 | Haplo | Yes | No | No | Steroid | No | NA | 45.8 | 5.4 | 6 | 0.2 | Poor/Poor | Moderna |
| P28 | F | 73 | AML | 14 | MMUD | No | Yes | Yes | Steroid, Tacrolimus | yes | Jakafi | 465 | 568.5 | 66 | 1.5 | Good/Good | Pfizer |
| V01 | M | ~35 | Healthy Control | | | | | | | | | | | | | | Moderna |
| V02 | F | ~55 | Healthy Control | | | | | | | | | | | | | | Moderna |
| V04 | M | ~80 | Healthy Control | | | | | | | | | | | | | | Moderna |

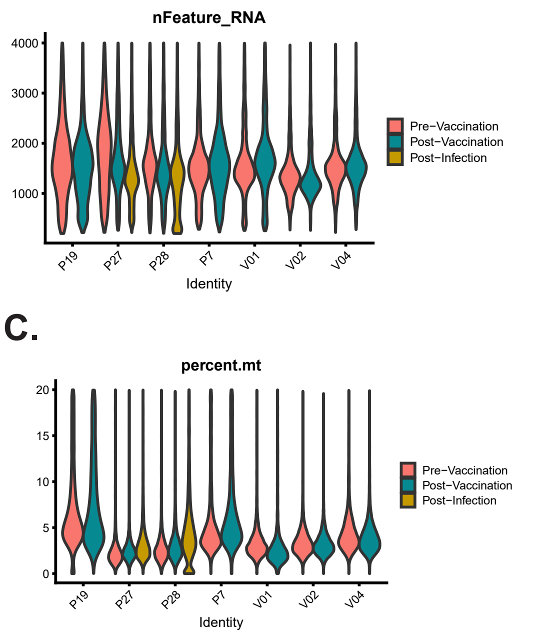

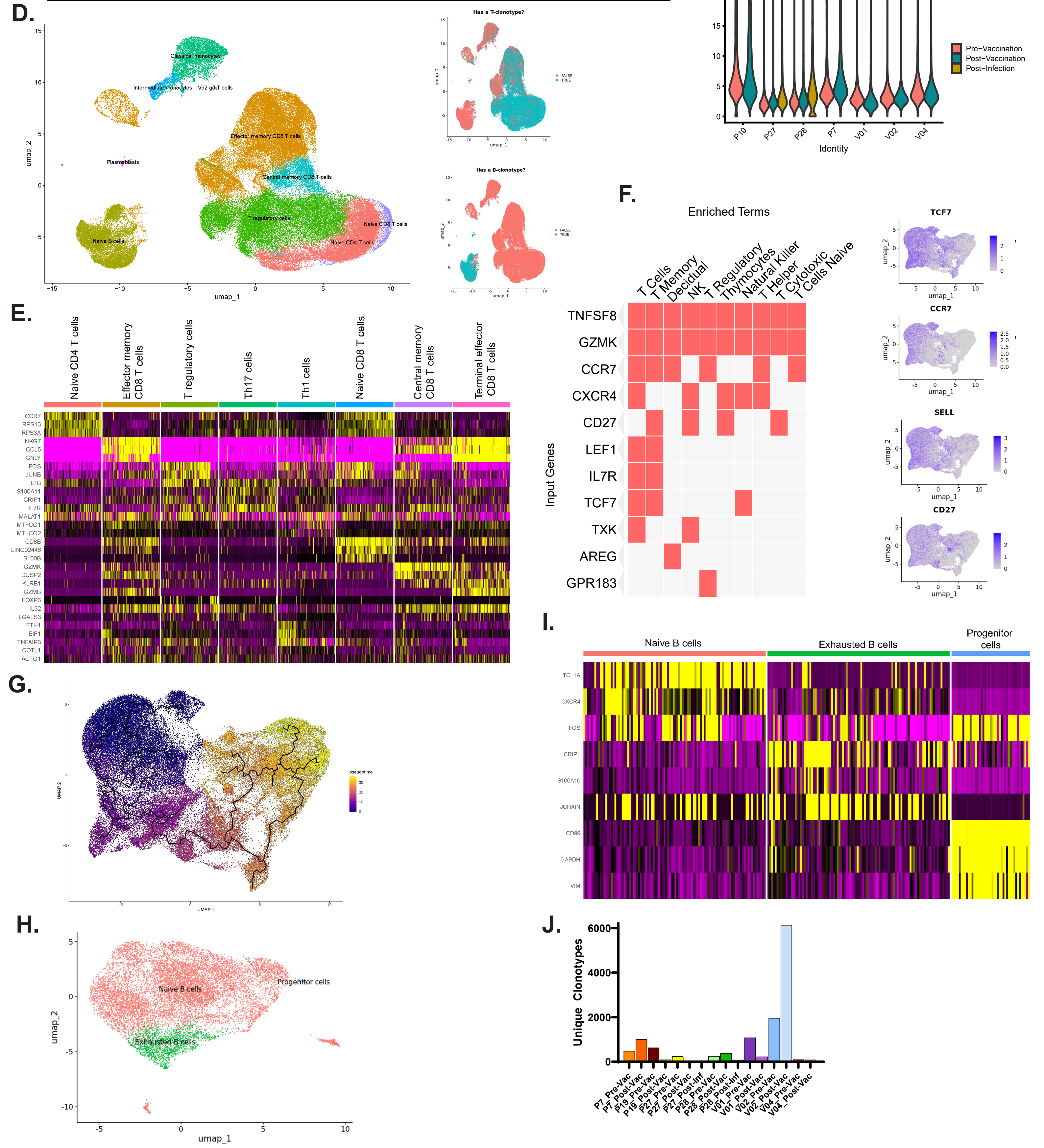

**Supplemental Figure 1**
